# *Pseudomonas aeruginosa* C-terminal processing protease CtpA assembles into a hexameric structure that requires activation by a spiral-shaped lipoprotein binding partner

**DOI:** 10.1101/2021.08.09.455718

**Authors:** Hao-Chi Hsu, Michelle Wang, Amanda Kovach, Andrew J. Darwin, Huilin Li

**Affiliations:** Department of Structural Biology, Van Andel Institute, Grand Rapids, Michigan, USA; Department of Microbiology, New York University Grossman School of Medicine, New York, New York, USA

## Abstract

*Pseudomonas aeruginosa* CtpA is a carboxyl terminal-processing protease that partners with the outer membrane lipoprotein LbcA to degrade at least five cell wall-associated proteins, four of which are cell wall hydrolases. This activity plays an important role in supporting *P. aeruginosa* virulence in a mouse model of acute pneumonia. However, almost nothing is known about the molecular mechanisms underlying CtpA and LbcA function. Here, we used structural analysis to show that CtpA alone assembles into an inactive hexamer comprising a trimer of dimers, which limits its substrate access and prevents nonspecific degradation. The adaptor protein LbcA is a right-handed open spiral with 11 tetratricopeptide repeats, which might wrap around a substrate to deliver it to CtpA for degradation. By structure-guided mutagenesis and functional assays, we also showed that the interfaces of the CtpA trimer-of-dimers, and an N-terminal helix of LbcA, are important for LbcA-mediated substrate degradation by CtpA both *in vitro* and *in vivo*. This work improves our understanding of the molecular mechanism of a CTP within the C-terminal processing peptidase-3 group.

**IMPORTANCE:** Carboxyl-terminal processing proteases (CTPs) are found in all three domains of life. In bacteria, some CTPs have been associated with virulence, raising the possibility that they could be theraputic targets. However, relatively little is known about their molecular mechanisms of action. In *Pseudomonas aeruginosa*, CtpA supports virulence by working in complex with the outer membrane lipoprotein LbcA to degrade cell wall hydrolases. Here, we report structure-function analyses of CtpA and LbcA, which reveals that CtpA assembles into an inactive hexamer comprising a trimer of dimers. LbcA is monomeric, with an N-terminal region important for binding to and activating CtpA, followed by a spiral structure composed of 11 tetratricopetide repeats, which could wrap around a substrate for delivery to CtpA. This work provides the first structure of a CTP-3 group member, revealing a unique mutimeric arrangement and insight into how this important proteolytic system functions.

## INTRODUCTION

A eubacterial cell is protected by a mesh-like cell wall of peptidoglycan (PG), which is composed of linear glycan strands with peptide side chains that cross-link to each other through peptide bonds (1). To accommodate growth, these cross-links must be cleaved so that nascent PGs can be inserted into the network (1–3). Several PG endopeptidases carry out this hydrolysis, including MepS, MepM, and MepH in *Escherichia coli* (4, 5). However, if not tightly regulated, their activity could lead to rupture of the PG sacculus and cell death. One way to control endopeptidase activity is through the relatively recently discovered carboxyl-terminal processing (CTP) protease system. CTPs belong to the S41 family of serine proteases (6). All CTPs have a PDZ domain — named because it was first noted in: <u>p</u>ostsynaptic density protein of 95 kDa, Drosophila <u>d</u>isc large tumor suppressor, and <u>z</u>onula occludens-1 protein — which plays roles in substrate recognition and protease regulation (7, 8). CTPs work within the cell envelope of Gram-negative and Gram-positive bacteria and have been linked to virulence (9–14).

The *E. coli* CTP Prc partners with the lipoprotein Nlpl to cleave the PG endopeptidase MepS (5). Prc is a bowl-shaped monomer, and the Nlpl adaptor forms a homodimer that binds to two separate molecules of Prc (15). In *Bacillus subtilis*, the CtpB protease processes and activates the intramembrane protease 4FA-4FB complex, thereby regulating spore formation (16, 17). CtpB has N-terminal and C-terminal dimerization domains, plus a cap domain preceding the protease core domain. Structural analysis revealed that CtpB assembles a dimeric self-compartmentalizing ring structure (18). The substrate peptide enters the proteolytic site via a narrow tunnel that is largely sequestered by the PDZ domain and becomes exposed only in the presence of a substrate. Therefore, the CtpB protease is reversibly activated by the substrate C-terminal peptide. In contrast to Prc and CtpB, *P. aeruginosa* CtpA has been assigned to the C-terminal processing peptidase-3 group (19). No structural studies have been reported so far for this group.

*Pseudomonas aeruginosa* is an opportunistic human pathogen. It is one of the leading causes of sepsis in intensive care units, and outbreaks of multidrug-resistant strains have been reported in hospitals (20–22). In contrast to *E. coli*, in which the only CTP present is Prc, *P. aeruginosa* has both the C-terminal processing peptidase-1 group member Prc, and the C-terminal processing peptidase-3 group member CtpA (10, 19). We reported previously that CtpA is required for normal function of the type 3 secretion system and for virulence in a mouse model of acute pneumonia (10). *P. aeruginosa* CtpA has 39% amino acid sequence identity with *B. subtilis* CtpB, which suggests that they might share a similar fold. However, CtpA has a much longer C-terminal region. This extended C-terminus might alter the oligomerization mode relative to that of CtpB (18). However, it is not known if or how CtpA assembles into a self-compartmentalizing structure to prevent nonspecific proteolysis. Unlike CtpB, CtpA is not directly activated by a protein substrate. Instead, it requires the adaptor protein LbcA, the <u>l</u>ipoprotein <u>b</u>inding partner of <u>C</u>tpA, for activity *in vivo* and *in vitro* (23).

LbcA is predicted to be an outer membrane lipoprotein, and to contain 11 tetratricopeptide repeats (TPRs). The TPR motif is a degenerate 34-amino-acid sequence that mediates protein-protein interactions (24–26). LbcA promotes CtpA protease activity, and *ctpA* and *lbcA* null mutants share common phenotypes, such as a defective type 3 secretion system and accelerated surface attachment (10, 23, 27). Five LbcA-CtpA substrates have been reported to date, and four of them are predicted to be PG cross-link hydrolases: the LytM/M23 family peptidases MepM and PA4404, and the NlpC/P60 family peptidases PA1198 and PA1199 (23, 27). Therefore, it appears that LbcA interacts with CtpA to assemble an active proteolytic complex, which controls the activity of these enzymes by degrading them. However, the molecular mechanisms underlying CtpA and LbcA function are unknown. Here, we describe structural and functional analyses of CtpA and LbcA. We show that CtpA alone assembles as an inactive trimer of dimers and that LbcA is a right-handed spiral that might wrap around a substrate protein. Structure-guided mutagenesis confirms the functional importance of CtpA interfaces and identifies the interface between CtpA and LbcA.

## RESULTS

### CtpA assembles a trimer-of-dimers hexamer in solution

CtpA is located in the periplasm, tethered to the outer membrane via its interaction with LbcA (23). Its N-terminal 23 residues are a type I signal sequence, which is followed by a 14-residue region that is enriched in alanine, glycine, and proline that contributes to a bacterial low complexity region and is predicted to be disordered (28). Therefore, we removed these 37 amino acids to produce a truncated CtpA protein for structural and functional studies (Supplemental Table 1, Fig. 1a). For simplicity, we refer to this ΔN37 construct as CtpA throughout the main text. The CtpA crystals diffracted X-rays to 3.5-Å resolution. We first used molecular replacement with the *B. subtilis* CtpB as a search model to solve the CtpA structure, but this did not lead to a satisfactory solution. We then produced selenomethionine-substituted CtpA crystals for SAD-based phasing. The derivatized crystals diffracted to 3.3 Å, leading to successful structural solution (Supplemental Table 2). CtpA formed a hexamer that comprised a trimer of dimers (Fig. 1b). This oligomer state is consistent with an estimated mass of a hexamer from the gel filtration profile (Fig. 1c). As expected, each CtpA protomer consists of an N-terminal dimerization region (NDR), a PDZ domain, a cap domain, a protease core domain, and a C-terminal dimerization region (CDR) (Fig. 1d). The PDZ domain is partially disordered, but we were able to generate a homolog PDZ model based on the CtpB structure and accurately docked the domain guided by two anomalous density peaks from selenomethionine residues 153 and 160 of the domain (Supplemental Figure 1). The loop connecting S10 and H6 (aa’s 378–411) within the CDR was disordered, and the last two residues (435-436) were not resolved.

**FIG 1.**
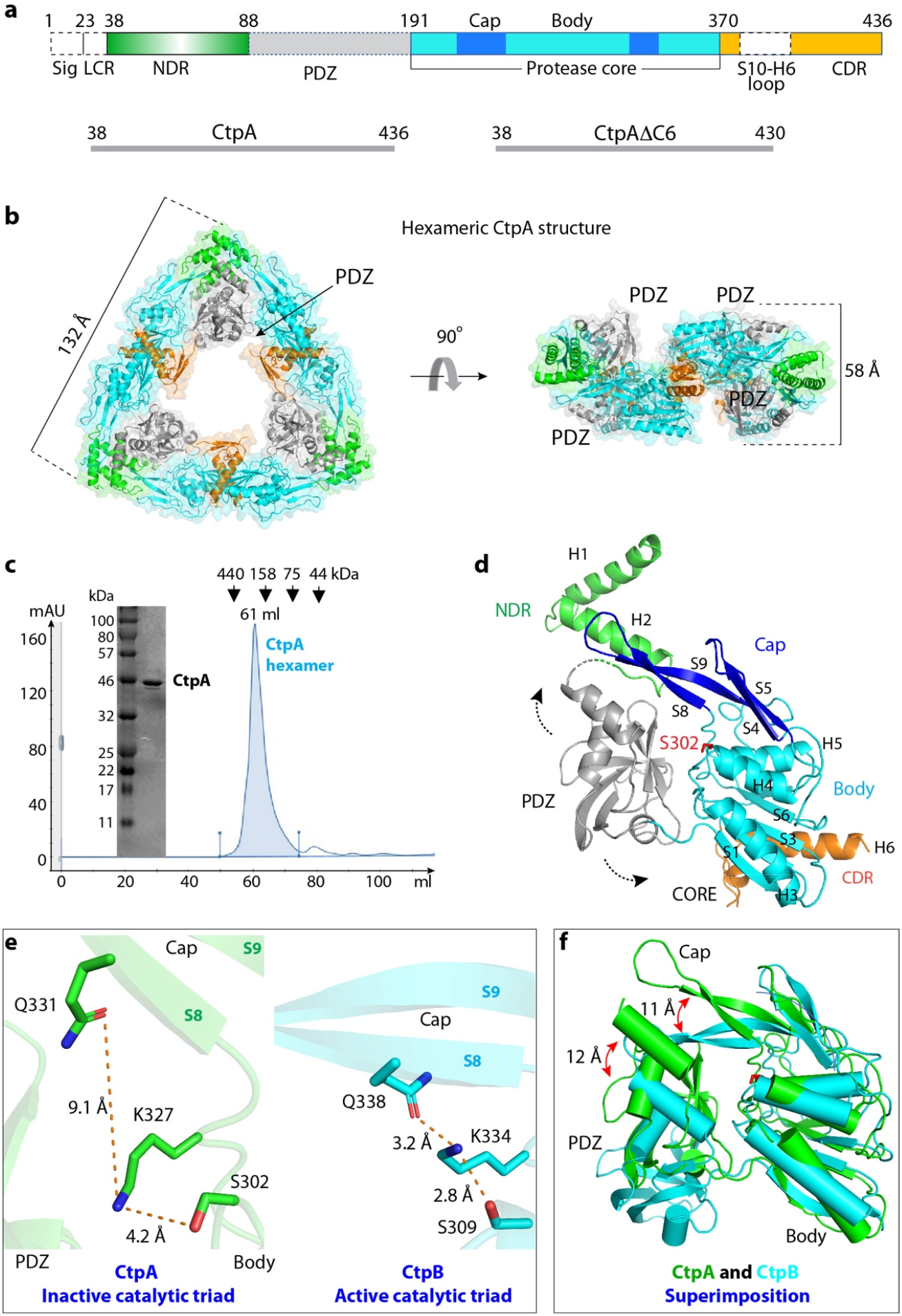
Overall structure of CtpA. (a) Top: CtpA domain organization. The dashed lines indicate the disordered regions (PDZ and the S10-H6 connecting loop) that are not resolved in the crystal structure. NDR, N-terminal dimerization region; CDR, C-terminal dimerization region. The sequence ranges of the two CtpA constructs used this study are shown in the lower panels. (b) Cartoon and transparent surface views of CtpA hexamer. The domains are colored according to the depiction in panel a. (c) SDS-PAGE analysis and gel filtration profile of CtpA). (d) A CtpA subunit in cartoon view. Secondary structural elements in CtpA are labeled, except in the PDZ domain. The two dashed arrows indicate the mobile PDZ domain in the CtpA hexamer. (e) Comparison of the catalytic triads of Pa CtpA and Bs CtpB (PDB ID 4C2E). The S8 and S9 labels refer to β-strands 8 and 9 in the cap region. (f) Superposition of the core domains of inactive Pa CtpA (green) and active Bs CtpB (cyan). Catalytic Ser-302 in CtpA and Ser-309 in CtpB are in red sticks. Red arrows indicate lifted-up (CtpA) and clamped-down positions (CtpB) of the cap subdomains.

### The CtpA hexamer alone in the absence of LbcA is in an inactive configuration

The CtpA core region contains the protease active site. There is a narrow tunnel dividing the core domain into the upper cap and lower body regions (Fig. 1d). In CtpB this tunnel was suggested to guide the substrate peptide into the proteolytic site (18). The cap of CtpA is a four-stranded β-sheet (S4, S5, S8, S9) located below the NDR. The body region is composed of a three-helix bundle (H3, H4, H5) and a five-stranded β-sheet (S1, S2, S3, S6, S7). The CtpA catalytic Ser-302 is located at the end of the narrow tunnel between the cap and the body region.

The CtpA hexamer is in an inactive conformation. The PDZ domain is in a position that blocks the narrow substrate peptide tunnel leading to Ser-302. Furthermore, the catalytic triad Ser-302–Lys-327–Gln-331 is well beyond hydrogen-bonding distance: Ser-302 and Lys-327 are 4.2 Å apart and Lys-327 and Gln-331 are 9.1 Å apart (Fig. 1e). We solved the crystal structure of CtpA that was catalytically inactive due to an S302A substitution and found that it was also in the inactive configuration (Supplemental Figure 2). For comparison, the catalytic triad Ser-309–Lys-334–Gln-338 in the protease-active *B. subtilis* CtpB are all within hydrogen-bonding distance (Fig. 1e). By superimposing the two proteases, we found that transition to the active form requires the CtpA cap domain to clamp down toward the catalytic site by 11 Å and also a large-scale movement of the associated PDZ domain by 12 Å (Fig. 1f, Supplemental Video 1). The CtpA structure is consistent with our previous observation that purified wild-type CtpA alone was inactive in degrading its substrates (23). The addition of purified LbcA was able to activate the protease of wild-type CtpA but not of the mutant CtpA(S302A). Therefore, unlike CtpB, which fluctuates between an active and inactive form in solution and can be activated by a protein substrate (18), the CtpA hexamer is locked in an inactive configuration and requires LbcA for activation (23).

### The C-terminal dimerization interface is important for full CtpA activity

*P. aeruginosa* CtpA dimerizes via its NDR in the same way as *B. subtilis* CtpB (Fig. 2a). However, the two CDRs of CtpA are far apart in this dimer, and six unhinged CDRs of three CtpA dimers interact to form the triangular trimer-of-dimer complex (Figs. 1b, 2a). In contrast, CtpB forms a parallel homodimer by both head-to-head N-terminal and tail-to-tail C-terminal interactions (Fig. 2b) (18). The CtpA NDR is composed of the H1 and H2 helices. The NDRs of two CtpA monomers form a domain-swapped, intermolecular 4-helix bundle, in which the NDR of one protomer reaches over to contact the cap domain of the other protomer (Fig. 2a, 2c). The four-helix bundle is at the vertex of the triangular complex. The NDR–NDR dimerization is driven by hydrophobic interactions. Specifically, Leu-69, Ala-73, and Met-77 are at the intersecting point of the two crossed H2’s. Met-77 hydrophobically interacts with Ala-73 and Leu-69 of the second protomer (Fig. 2c). H1 of protomer 1 is nearly parallel to H2 of protomer 2, with the H1 residues Leu-46, Phe-49, Val-52, Leu-53, and Val-56 hydrophobically interacting with the H2 residues Leu-70, Ile-74, Met-77, Leu-78, and Leu-81. The Leu-46 and Phe-49 of the two H1’s are also within the van der Waals distance. The hydrophobic NDR–NDR interaction of the CtpA dimer resembles that in the CtpB dimer, consistent with the conserved sequence in this region (Fig. 2e).

**FIG 2.**
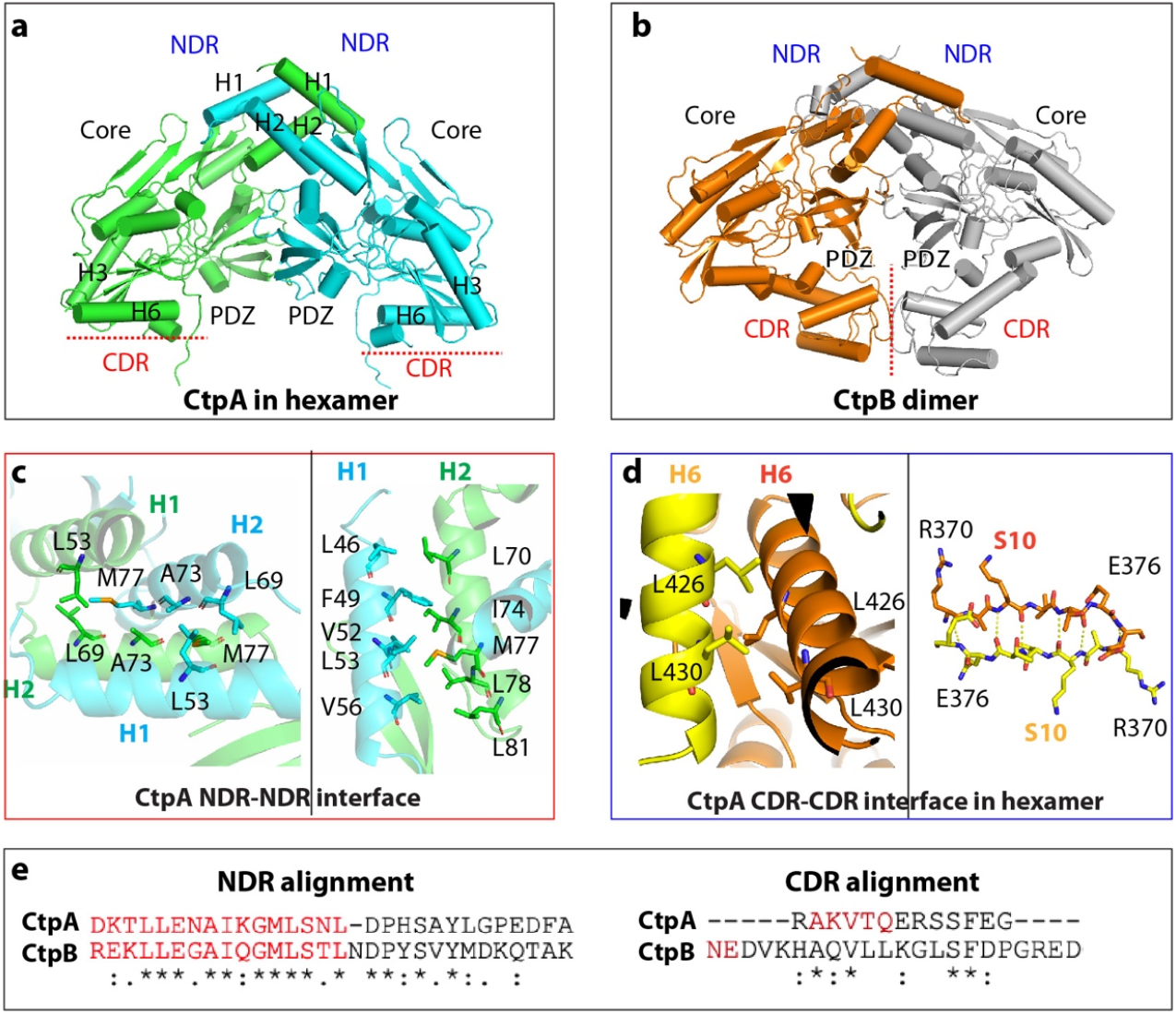
Different oligomerization modes of Pa CtpA and Bs CtpB. (a) A CtpA dimer extracted from the CtpA hexamer. (b) Bs CtpB dimer (PDB ID 4C2E). (c) The N-terminal dimerization interface of CtpA involves hydrophobic interactions between two H2 (left) and between H1 and H2 (right). (d) The C-terminal dimerization interface of CtpA involves a short leucine zipper-like interaction between two H6 helices and antiparallel β-sheet formation between two S10. (e) Alignment of the conserved NDR sequence and divergent CDR between Pa CtpA and Bs CtpB.

The C-terminal dimerization interface of CtpA involves the β-strand S10 and the helix H6 (Fig. 2d). The two S10 β-strands (Arg-370 to Glu-376) form an intermolecular, antiparallel β-sheet that reinforces the dimer interface. The two H6’s (Tyr-418 to Gly-435) are orthogonal to each other but form a short leucine zipper in the middle section mediated by Leu-426 and Leu-430. The CtpA C-terminal dimer interface is not like that of CtpB, which dimerizes via interaction in a different region. Indeed, the C-terminal sequence of *P. aeruginosa* CtpA is not conserved in *B. subtilis* CtpB (Fig. 2e), which is consistent with their different oligomeric assembly. To investigate the functional importance of the unique C-terminal dimerization interface of CtpA, we constructed a mutant with a partially disrupted H6 helix by removing the last six residues Ser-431 to Asn-436; CtpA(ΔC6). We found that CtpA(ΔC6) eluted from a gel filtration column as a dimer (Fig. 3a), so the C-terminal truncation prevented CtpA hexamer formation.

**Fig. 3.**
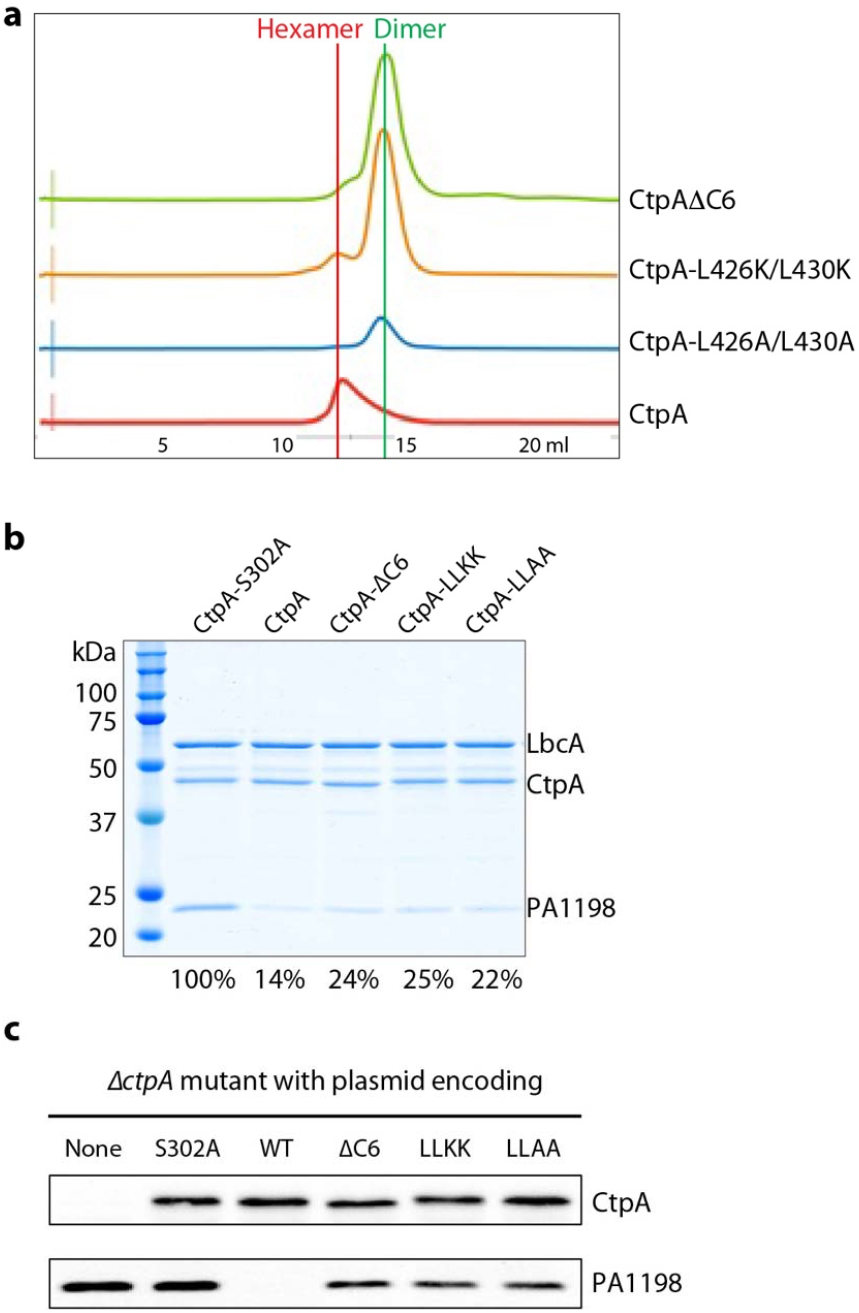
Full protease activity of CtpA requires C-terminal dimerization region. **a**) Elution profiles of wild-type and mutant CtpA proteins. **b**) Substrate degradation assay *in vitro*. His_6_-PA1198 served as the substrate for the assay. Gels from a single experiment are shown, but the amount of PA1198 degradation is the average from two independent experiments, determined as described in Methods. The number below each lane is the percentage of remaining PA1198 after 3 h, relative to the first lane using the inactive CtpA. **c**) Substrate degradation *in vivo*. Plasmid-encoded protease-dead CtpA(S302A), wild-type (WT), ΔC6, L426K/L430K (LLKK), L426A/L430A (LLAA) were produced in a *P. aeruginosa* Δ*ctpA* strain. None = empty plasmid vector control. The CtpA proteins and accumulation of the PA1198 substrate were detected by immunoblot analysis with polyclonal antisera.

We next asked whether the C-terminal dimerization interface, present in the intact CtpA hexamer but disrupted in the CtpA(ΔC6) dimer, is required for normal CtpA function. In an *in vitro* assay, we found that the CtpA(ΔC6) dimer was less active than the CtpA hexamer in degrading the model substrate PA1198, in reactions that also contained LbcA (compare lanes 3 and 4 in Fig. 3b).

To further investigate the functional significance of the leucine zipper in H6, we generated two CtpA double-mutant constructs with L426K/L430K and L426A/L430A substitutions that disrupted the leucine zipper interaction. The two purified mutant proteins were mainly dimeric and failed to assemble into a hexamer, based on their gel filtration profiles (Fig. 3a). Their LbcA-activated *in vitro* protease activity toward PA1198 was lower relative to wild-type CtpA (Fig. 3b, compare lanes 5 and 6 to lane 3). However, like the CtpA(ΔC6) mutant, the leucine zipper mutants retained significant protease activity relative to the protease dead mutant CtpA(S302A) (Fig. 3b, compare lanes 5 and 6 with lane 2).

We extended our analysis by constructing plasmids encoding derivatives of full-length CtpA with the ΔC6, L426K/L430K (LLKK), or L426A/L430A (LLAA) mutations. After introducing these plasmids into a *P. aeruginosa ΔctpA* mutant, immunoblot analysis showed that the steady-state levels of the mutant proteins were similar to the wild-type (Fig. 3c). Therefore, the mutations did not affect CtpA stability *in vivo*. However, PA1198 accumulated in the presence of these three mutants relative to the wild-type, although not as much as in the presence of CtpA(S302A) (Fig. 3c). This suggests that the protease activity of the three mutants was reduced, but not abolished, which is consistent with the *in vitro* analysis (Fig. 3b). Therefore, our studies suggest that the hexameric assembly is important for proper CtpA function.

### LbcA forms a spiral that could wrap around CtpA substrates

During the maturation of the lipoprotein LbcA, its N-terminal 16 residues are removed, exposing Cys-17 (Fig. 4a). Then Cys-17 is lipidated so that the N-terminus can be anchored in the outer membrane from the periplasmic side (29–31). LbcA is able to bind to CtpA and its substrates independently (27). Therefore, to begin to understand how LbcA might be capable of these separate interactions, we tried to solve the LbcA crystal structure. We were able to crystallize LbcA with its N-terminal 48 residues removed (ΔN48). The crystal structure was solved to 3.5-Å resolution by the SAD method with selenomethionine-derivatized LbcA(ΔN48) crystals (Fig. 4a-d). This structure contains only α-helices and connecting loops, with a total of 29 α-helices. We resolved all 11 predicted tetratricopeptide repeats, comprising 22 α-helices from helix-7 (H7) to H28. The sequences of the 34-residue TPRs are largely conserved (Fig. 4b). Gly/Ala at position 8 and Ala at positions 20 and 27 are most conserved in the TPR family, although none of these are invariant. Interestingly, TPR1, TPR5, and TPR10 contain only 33 amino acids, but they all contain the signature residues Gly/Ala at position 8 and Ala at position 20, and TPR1 and TPR10 also contain a signature Leu at position 24. Within each TPR, the first helix (TPR-A) lines the inner surface and the second helix (TPR-B) lines the outer surface of the ring structure. Alignment of the 11 TPRs in LbcA(ΔN48) showed that more-conserved hydrophobic residues are concentrated in the TPR-A helix and more-conserved charged residues are distributed in TPR-B helix and in the turns connecting TPR-A and TPR-B, suggesting that the inner and the outer surfaces of the ring may have distinct functions.

**FIG 4.**
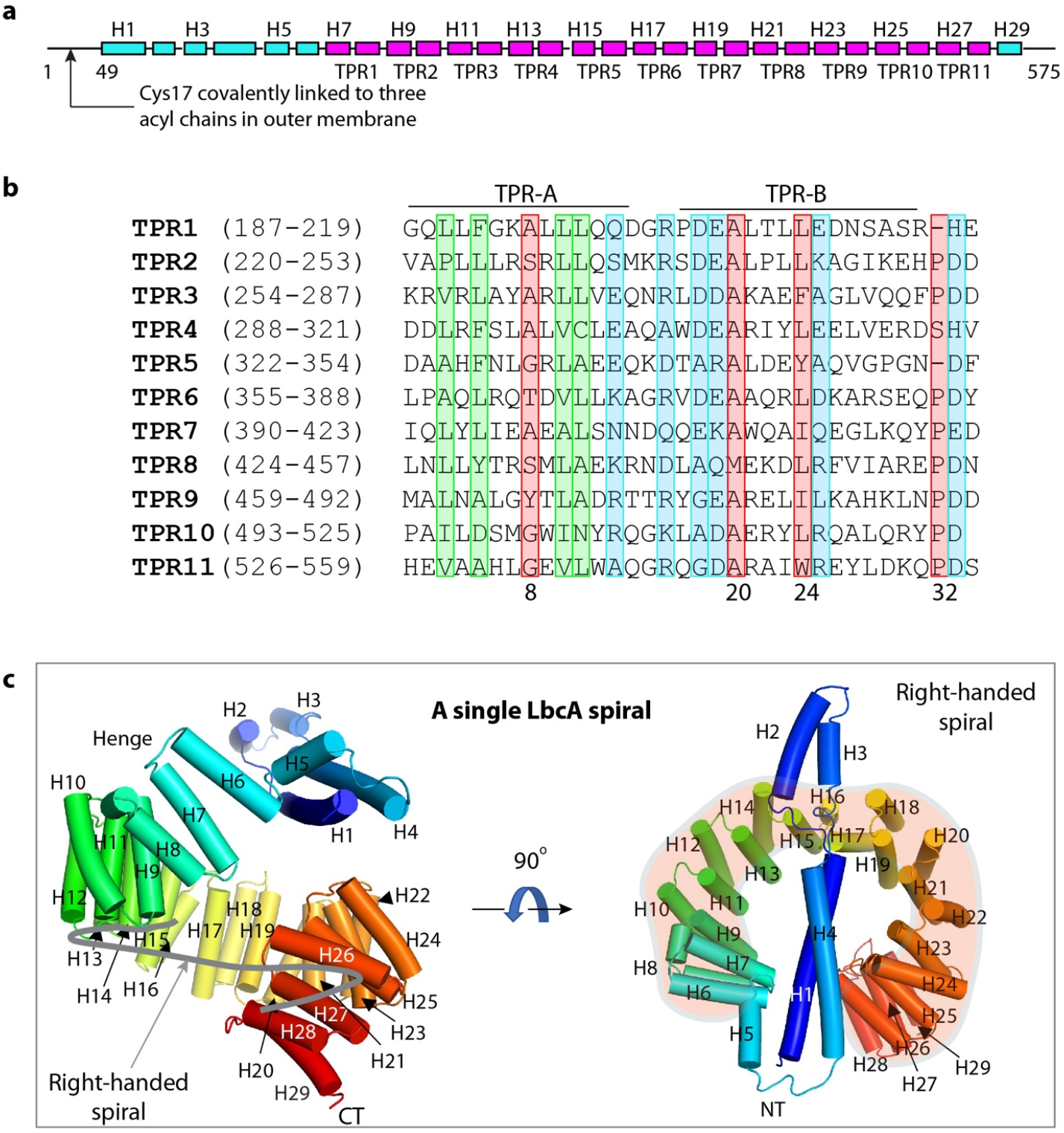
Overall structure of LbcA. (a) Domain organization of LbcA. The TPR motifs are shown in magenta. Also shown are two LbcA constructs used in this study. (b) Sequence alignment of the 11 TPRs of LbcA. The signature residues of TPR are marked in the red rectangles. The highly homologous hydrophobic residues are in green boxes and the highly homologous charged residues are in the blue boxes. (c) The crystal structure of LbcA(ΔN48) contains an N-terminal extension with 4 helices and a C-terminal superhelix composed of 11 TPRs. The thick gray curve in the left panel follows the right-handed spiral feature of the TPRs.

The 11 TPRs form a notched ring, with an outer diameter of about 6 nm and an inner diameter of about 3 nm (Fig. 4c). It is a right-handed spiral structure, because the first TPR is slightly above, and the last TPR is below the ring. The first 4 α-helices at the N terminus form an elongated extension that partially caps the TPR ring. Helices H5 and H6 serve as a hinge that links the N-terminal helical extension and the TPR spiral. However, the loop connecting H4 and H5 (aa’s 163-171) has a high crystallographic B-factor (150-200 Å^2^) that is indicative of flexibility. Therefore, this loop and the H5-H6 hinge are likely to allow relative motion of the NT helical extension with respect to the TPR spiral. The TPR ring is capped at the end by the single short α-helix, H29.

LbcA(ΔN48) purified as a monomer in solution (Fig. 5a-b). However, four LbcA(ΔN48) molecules formed an interlocked tetramer as a dimer of dimers in the crystal (Fig. 5c-d). The chamber of the TPR spiral of the first LbcA is occupied by the H3-H4 of a second LbcA molecule on the top and by the H1-H2 of a third LbcA molecule on the side (Fig. 5d). Therefore, there is a 4-helix bundle inside the first LbcA TPR spiral. We suggest that this bundle may mimic a substrate and that the LbcA spiral may wrap around a substrate to target it to CtpA for degradation (Fig. 5e). In this scenario, the conserved and hydrophobic residues of the TPR-A helices lining the inner surface of the TPR spiral may participate in binding that substrate.

**FIG 5.**
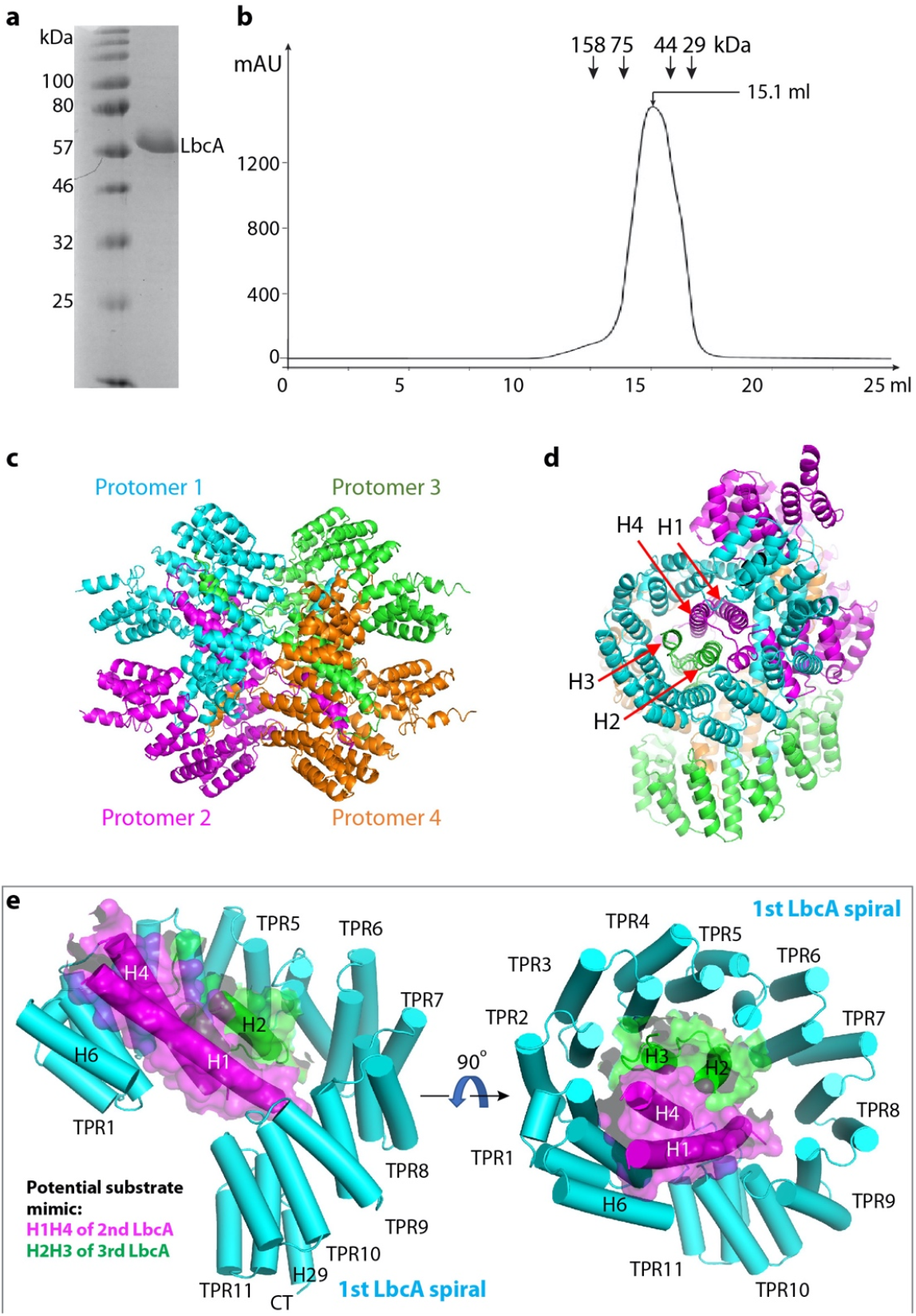
LbcA is a monomer in solution but assembles a tetramer in crystal. (a) Coomassie blue stained SDS-PAGE gel of the purified of LbcAΔN48. (b) Superdex-200 elution profile of LbcAΔN48. LbcA was eluted from gel filtration column at a volume corresponding to the monomeric state. (c) LbcA forms an intertwined tetramer in the crystal lattice. The four protomers are individually colored. (d) This LbcA tetramer view shows that the helices H1 and H4 of protomer 2 and helices H2 and H3 of protomer 3 form a 4-helix bundle inside the super helical coil of protomer 1 in the crystal lattice. (e) H2H3 of a second LbcA (magenta cartoon and transparent surface) and H1H4 of a third LbcA (light blue cartoon and transparent surface) insert into the first LbcA spiral, likely mimicking the substrate binding by the first LbcA spiral.

### The LbcA H1 is essential for binding to and activating the CtpA protease

To identify the binding interface between CtpA and LbcA, we produced a series of N- and/or C-terminal-truncated LbcA mutants and did a CtpA pulldown assay (Fig. 6a). We found that removing the C-terminal five residues of LbcA did not affect the LbcA–CtpA interaction. All constructs containing the complete H1-H29 region pulled down CtpA. However, LbcA with either H1 truncation (His_6_-LbcAΔN84) or H1-H4 truncation (His_6_-LbcAΔN165) failed to pull down CtpA. This result suggests that the LbcA NT helical extension, in particular H1, is required for LbcA binding to CtpA, and that LbcA H1 directly participates in the binding.

**FIG 6.**
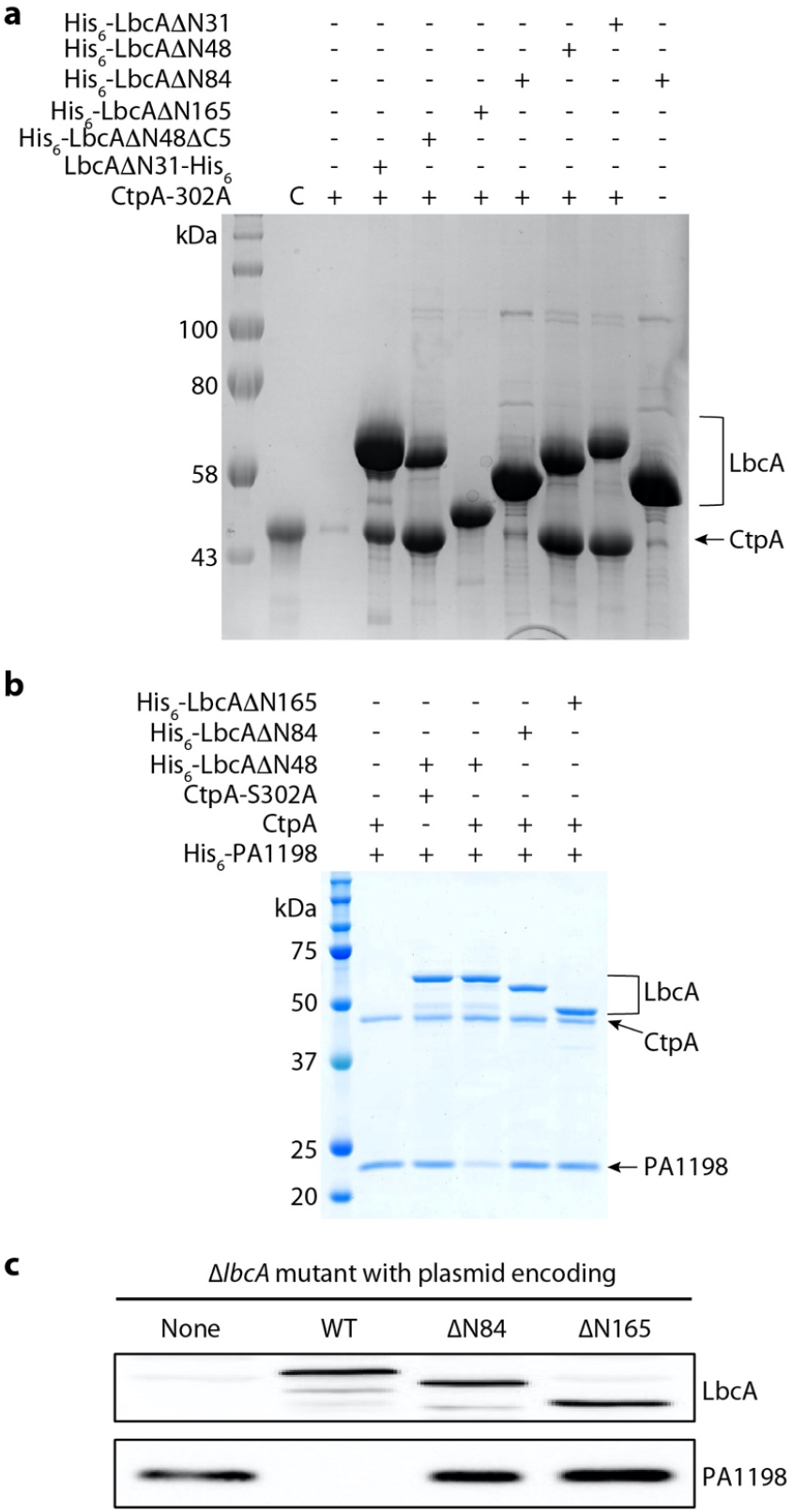
The LbcA N-terminal extension is essential for the protease activity of CtpA. (a) CtpA pull-down assay using various N-terminal and C-terminal deletion mutants of LbcA. Lane 2 is the CtpA input control (C). Lane 3 is the background binding of CtpA to Ni beads. (b) *In vitro* substrate (PA1198) degradation assay. (c) Substrate degradation *in vivo*. Plasmids encoding wild-type LbcA (WT), LbcA(ΔN84), or LbcA(ΔN165) were transformed into a *P. aeruginosa ΔlbcA* mutant. None = empty plasmid vector control. The LbcA proteins and accumulation of the PA1198 substrate were detected by immunoblot analysis with polyclonal antisera.

We then carried out both *in vitro* and *in vivo* assays to monitor degradation of the PA1198 substrate (23). We first incubated purified CtpA with separately purified LbcA proteins and PA1198. As expected, PA1198 was degraded in the presence of His_6_-LbcAΔN48, which had the intact NT helical extension (Fig. 6b). However, PA1198 was not degraded in the presence of His_6_-LbcA(ΔN84) (missing H1) or His_6_-LbcA(ΔN165) (missing H1-H4). In fact, the outcomes of reactions with these two truncations were indistinguishable from those containing no LbcA, or those using the protease dead CtpA(S302) (Fig. 6b). In a Δ*lbcA* mutant *P. aeruginosa* strain, plasmids encoding LbcA with in frame deletions equivalent to the ΔN84 or ΔN165 constructs failed to activate CtpA, as revealed by the accumulation of PA1198, which was present at a similar level as in a strain without any LbcA (Fig. 6c). Therefore, both pulldowns and *in vitro* and *in vivo* activity assays pinpointed the LbcA H1 as a key binding element for activation of the CtpA protease.

## DISCUSSION

The LbcA–CtpA system supports the function of the *P. aeruginosa* type 3 secretion system, is required for virulence in a mouse model of acute infection, and affects surface attachment (10, 23). Furthermore, four CtpA substrates are PG cross-link hydrolases, which means that the LbcA–CtpA system affects the integrity of a crucial cell envelope component and perhaps the most important antibiotic target, the cell wall. Therefore, this system could be an effective antibiotic target, and the structural analysis reported here may aid the development of such antibiotics. Here, we have shown that CtpA assembles as a hexamer. However, the hexamer alone is inactive, because the catalytic triad Ser302–Lys327–Gln331 is not in hydrogen-bonding distance. We also found that the CtpA partner protein LbcA has an N-terminal helical region that is needed to bind to CtpA, and a large spiral cavity that has the capacity to wrap around a substrate for delivery to CtpA.

How the interaction with LbcA converts CtpA into an active protease is currently unclear. Elucidation of the activation mechanism requires the determination of the CtpA-LbcA complex. However, our previous experiments suggested that most, if not all, CtpA in the cell is bound to LbcA (23). CtpA fractionates with the membrane fraction in *lbcA*^+^ cells, but is in the soluble periplasmic fraction in Δ*lbcA* cells, and when CtpA or LbcA is purified from *P. aeruginosa*, they copurify with a lot of the other one (23). Also, the pulldown assays done here show that LbcA and CtpA can interact in the absence of substrate (Fig. 6a). Therefore, it is possible that the LbcA-CtpA complex fluctuates between inactive and active states, and perhaps the presence of a substrate would stabilize the enzyme-adaptor-substrate in the active state for productive substrate hydrolase degradation.

The PDZ domain is the C-terminal peptide substrate-binding element of CTPs (32). The ability of PDZ domains to move in order to accommodate the incoming C-terminal substrate peptide partially accounts for the substrate delivery–based activation mechanism of the CTPs. The PDZ domain is the most flexible region in the CtpB structure and acts as an inhibitor by blocking the substrate peptide binding in the inactive form, but moves away to form a narrow tunnel for a substrate peptide in the active form (18). Similarly, the PDZ domain of CtpA is highly flexible and largely invisible in the crystal structure of the inactive CtpA hexamer, suggesting a CtpA activation mechanism similar to that of CtpB. In the case of Prc from *E. coli*, the PDZ domain acts as an activator rather than an inhibitor, but its movement is still a key feature of the activation mechanism (8).

*P. aeruginosa* CtpA requires the partner lipoprotein LbcA for activation and targeted proteolysis (23). This is a variation in the general substrate activation mechanism. In this regard, the LbcA–CtpA system is analogous to the *E. coli* NlpI–Prc system, in which the TPR-containing adaptor NlpI plays the dual function of delivering the PG hydrolase MepS to the Prc protease for degradation, and activating the protease (8, 15). Despite this functional similarity, significant differences exist between these two systems. The two proteases are in different C-terminal processing peptidase families; *E. coli* Prc is much larger than *P. aeruginosa* CtpA; and Prc functions as a monomer, not a hexamer. Therefore, the Prc N-terminal and C-terminal helical domains are not involved in oligomerization; instead, they wrap around the protease core, perhaps to limit substrate access to the protease. Nlpl has a total of 14 helices with four TPRs, compared with the 29 helices and 11 TPRs of LbcA. Although NlpI and LbcA both contain TPRs, the primary sequences of the proteins are not similar. Finally, NIpI functions as a dimer, unlike the monomeric state of LbcA.

The LbcA–CtpA system contributes to the virulence of *P. aeruginosa*. This makes understanding the structure and function of this proteolytic system broadly significant. Future challenges include understanding exactly how LbcA activates CtpA, and how protein substrates, not just peptides, are specifically recognized by the multitude of CTPs and/or their adaptor proteins for tightly regulated proteolysis.

## MATERIALS AND METHODS

### Purification of CtpA

DNA encoding amino acid 38 to the C-terminus of CtpA was sucloned into plasmid pET15b between the NdeI and XhoI sites. The plasmid encoded N-terminal His_6_-tagged CtpA(ΔN37). Similar plasmids encoding CtpA(ΔC6), CtpA-L426K L430K or CtpA-L426A L430A were constructed using reverse primers that incoprporated the C-terminal mutations. *E. coli* BL21(DE3) transformants were grown at 37°C to OD_600_ = 0.6-0.7 before being induced with 0.5 mM IPTG and incubated at 16°C overnight. Cells were harvested by centrifugation and lysed by passing through a microfluidizer cell disruptor in 10 mM potassium phosphate, pH 8.0, 10 mM imidazole, 0.25 M NaCl. The homogenate was clarified by centrifuging at 27,000 x *g* and the supernatant was applied to a HiTrap-Ni column (GE Healthcare) pre-equilibrated with the lysis buffer. Proteins were eluted with a 10–300 mM imidazole gradient in 10 mM potassium phosphate, pH 8.0, containing 0.25 M NaCl. Fractions containing His_6_-CtpA were collected. The N-terminal His tag was removed using thrombin (0.5 units/mg) by dialyzing against 20 mM Tris, pH 8.0, 150 mM NaCl overnight at 4 °C. Untagged CtpA was further purified with HiTrap-Q in 10 mM Tris, pH 8.0, and a 50–500 mM NaCl gradient and polished by gel filtration in 10 mM Tris, pH 8.0, and 150 mM NaCl using Superdex 200 prep grade column (16 x 600 mm, GE Healthcare).

### Purification of LbcA

DNA encoding amino acid 49 to the C-terminus of LbcA (LbcA[ΔN48]) was subcloned into the NdeI and HindIII sites of plasmid pET24b. Plasmids encoding N-terminal His_6_-tagged LbcA proteins were constructed similarly by subcloning fragments into the NdeI and BamHI sites of plasmid pET15b. *E. coli* BL21(DE3) transformants were grown at 37°C to OD_600_ = 0.5 before being induced with 0.5 mM IPTG and incubated at 37°C for another 3 h. His_6_-tagged LbcA protein was purified with HiTrap-Ni in 10 mM potassium phosphate, pH 8.0, 0.25 M NaCl, and a 10– 300 mM imidazole gradient, followed by HiTrap-Q in 10 mM Tris, pH 8.0, and a 50–500 mM NaCl gradient. The final polish of LbcA was done in a Superdex 200 prep-grade column preequilibrated with 10 mM Tris, pH 8.0, and 150 mM NaCl.

### Protein crystallization and structural solution

CtpA(ΔN37) was crystallized at 20°C by the sitting-drop vapor diffusion method using 0.1 M sodium acetate, pH 4.0, and 0.6 M ammonium dihydrogen phosphate at a concentration of 33 mg/mL. Diffraction data were collected at the Lilly Research Laboratories Collaborative Access Team (LRL-CAT) beamline of the Advanced Photon Source (APS) at Argonne National Laboratory and processed with Mosflm software. Se-derived crystals diffracted to 3.3 Å, which was better than the native crystals, so the Se-derived data were used to solve the CtpA structure. Autosol of Phenix was used to locate Se sites and the resulting map was used to build the initial model. However, the map quality from the SAD method was not good enough to build the CtpA model due to a low anomalous signal (only 4 high-occupancy Se were identified in one asymmetric unit). In combination with molecular replacement, the structure of CtpA(ΔN37) was determined by combining MR-SAD using the core domain of *B. subtilis* CtpB (PDB ID 4C2C) as the search model. The structure of CtpA(ΔN37, S302A) was solved by PHASER of Phenix using CtpA(ΔN37) as the search model.

LbcA(ΔN48) was crystallized at 20°C by the sitting-drop vapor diffusion method using 0.1 M sodium acetate, pH 4.6, and 1.9 M ammonium dihydrogen phosphate at a concentration of 45 mg/mL. Diffraction data were collected at the Life Sciences Collaborative Access Team (LS-CAT) beamline of APS and were processed with Mosflm. The best data set was from Se-derived crystals which diffracted to 3.5 Å, so the Se-derived diffraction data were used to solve the LbcA structure. Se sites were determined by the SAD method using Autosol of Phenix (33). All models of CtpA and LbcA were built in Coot (34) and refined with Phenix.refine (35).

The crystal structures of CtpA, CtpA(S302A), and LbcA have been deposited in the Protein Data Bank under accession codes 7RPQ, 7RQH, and 7RQF, respectively.

### *In vitro* proteolysis assay

CtpA and LbcA proteins were purified as described above. To purify His_6_-PA1198, *E. coli* strain M15 [pREP4] (Qiagen) containing pAJD2948 was grown in 500 mL of LB broth at 37°C to an OD_600_ of 0.6-1.0. Protein production was induced by adding 1 mM IPTG and incubation at 37 °C for 3 h, after which cells were harvested by centrifugation. His_6_-PA1198 was purified under native conditions by nickel–nitrilotriacetic acid–agarose affinity chromatography in buffer containing 50 mM NaH_2_PO_4_ and 300 mM NaCl, as recommended by the manufacturer (Qiagen). Protein was eluted in 50 mM NaH_2_PO_4_, 300 mM NaCl, 50 mM imidazole, pH 8. Assays were done as described previously (23, 36). Reactions were terminated by adding SDS-PAGE sample buffer and incubating at 90°C for 10 min. Samples were separated by SDS-PAGE and stained with ProtoBlue Safe (National Diagnostics). For experiments with mutant CtpA proteins, ImageJ analysis was used to determine the densities of the His_6_-PA1198 bands. The amount of His_6_-PA1198 degraded in each reaction was determined by comparing His_6_-PA1198 band density to that in the reaction with inactive CtpA(S302A). Average values from two independent experiments are reported.

### *In vivo* CtpA activity assay

Wild-type *ctpA* was amplified from *P. aeruginosa* PAK chromosomal DNA using a primer that annealed about 30 bp upstream of the *ctpA* start codon and a reverse primer that annealed immediately downstream of the stop codon. A fragment encoding CtpA(ΔC6) was generated with a reverse primer that annealed downstream of codon 430 and incorporated a stop codon. Fragments encoding CtpA-L426K L430K or CtpA-L426A L430A were generated with reverse primers that incorporated the mutagenic mismatches. A fragment encoding CtpA(S302A) was generated by amplifying *ctpA* from *P. aeruginosa* strain AJDP1140 DNA using the same primer pairs used to amplify wild-type *ctpA*. All fragments were cloned into pHERD26T using EcoRI and XbaI restriction sites added by the amplification primers.

Fragments encoding LbcA(ΔN84) and LbcA(ΔN165) were generated by amplifying one fragment with a forward primer that annealed ~ 40 bp upstream of the *lbcA* start codon and a reverse primer that annealed at codon 31, and a second fragment with a forward primer that annealed at codon 85 or codon 166 and a reverse primer that annealed at the *lbcA* stop codon. The forward primer for the second fragments had a 5’ tail complementary to the end of the first fragment (codons 25-31). The first fragment and second fragments were joined by splicing overlap extension PCR (37). The resulting fragments encoded LbcA with internal deletions that removed codons 32-84 (ΔN84) or codons 32-165 (ΔN165). Codons 1-31 were retained, encoding the signal sequence followed by 15 amino acids corresponding to the N-terminus of mature wild-type LbcA, to ensure normal signal sequence processing, lipidation, and outer membrane trafficking. A fragment encoding wild-type *lbcA* was generated by amplifying *lbcA* from *P. aeruginosa* DNA using a forward primer that annealed ~ 40 bp upstream of the start codon and a reverse primer that annealed at the *lbcA* stop codon. All fragments were cloned into pHERD26T using EcoRI and HindIII restriction sites added by the amplification primers.

Plasmids were introduced into Δ*ctpA* or Δ*lbcA* mutants by electroporation (38). Saturated cultures were diluted into 5 mL of LB broth containing 75 μg/mL tetracycline, in 18-mm diameter test tubes, at OD_600_ of 0.05. Cultures were grown on a roller drum at 37 °C for 5 h. Cells were harvested by centrifugation and resuspended in SDS-PAGE sample buffer at equal concentrations based on culture OD_600_. Samples were separated by SDS-PAGE and transferred to nitrocellulose by semi-dry electroblotting. Chemiluminescent detection followed incubation with polyclonal antisera against CtpA, LbcA, or PA1198, and then goat anti-rabbit IgG (Sigma) horseradish peroxidase conjugate.

## ACKNOWLEDGEMENTS

Diffraction data for CtpA were collected at the Lilly Research Laboratories Collaborative Access Team (LRL-CAT) beamline and diffraction data of LbcA were collected at the Life Sciences Collaborative Access Team (LS-CAT) beamline of the Advanced Photon Source (APS). The APS is a U.S. Department of Energy (DOE) Office of Science User Facility operated for the DOE Office of Science by Argonne National Laboratory under Contract No. DE-AC02-06CH11357. Cryo-EM data were collected at the David Van Andel Advanced Cryo-Electron Microscopy Suite at Van Andel Institute. We thank Dolonchapa Chakraborty for supplying plasmid pAJD2948 and His_6_-PA1198 protein, Gongpu Zhao and Xing Meng for help with data collection, and David Nadziejka for technical editing of the manuscript. This study was supported by U.S. National Institutes of Health grant R01 AI136901 (to A.D.) and by Van Andel Institute (to H.L.). The content is solely the responsibility of the authors and does not necessarily represent the official views of the National Institutes of Health.

---

**Supplemental Table 1.**
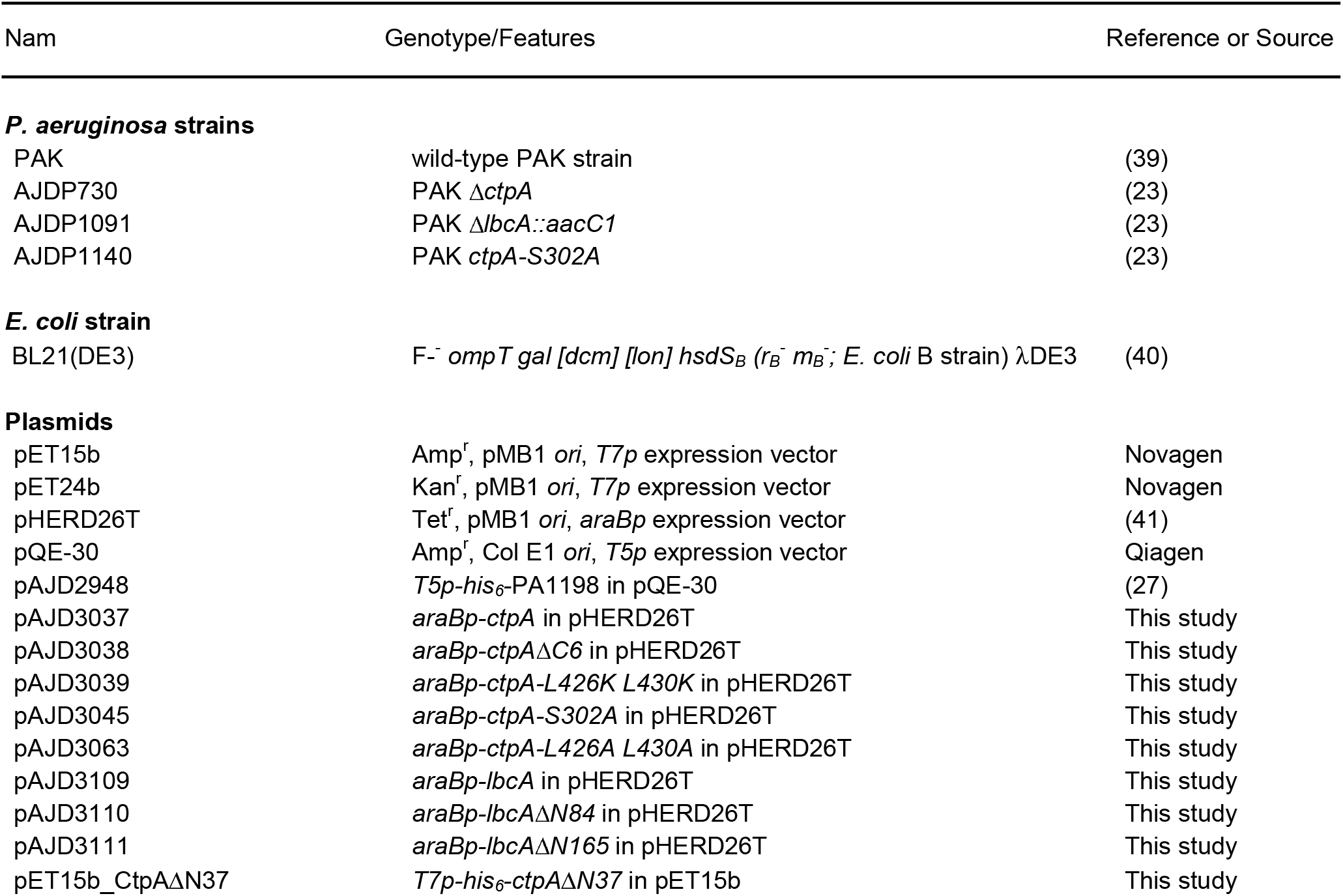
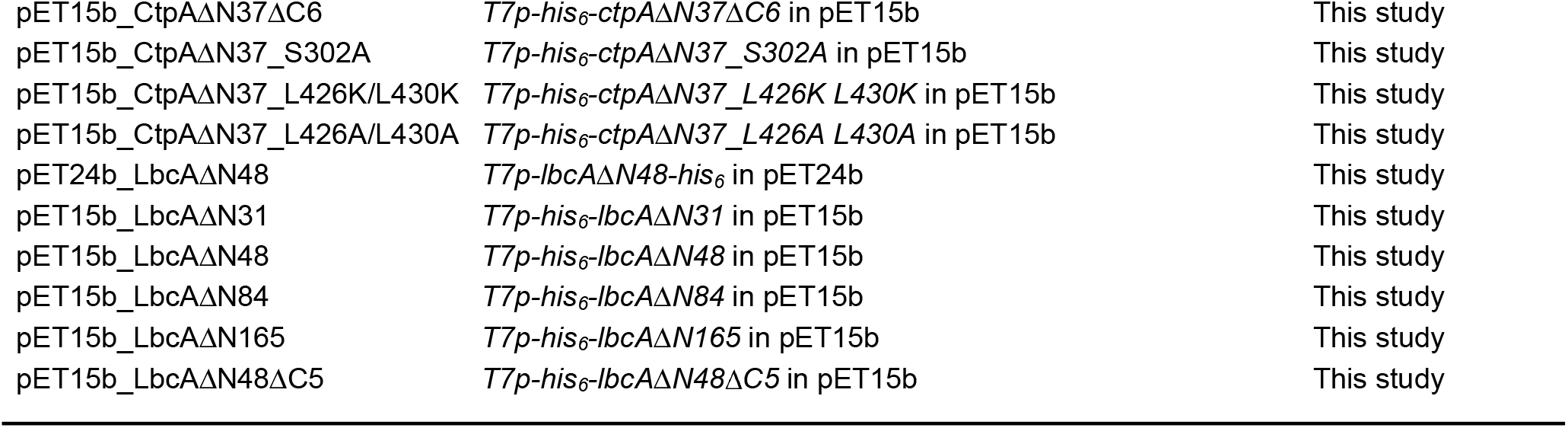
Strains and plasmids used in this study.

**Supplemental Table 2.**
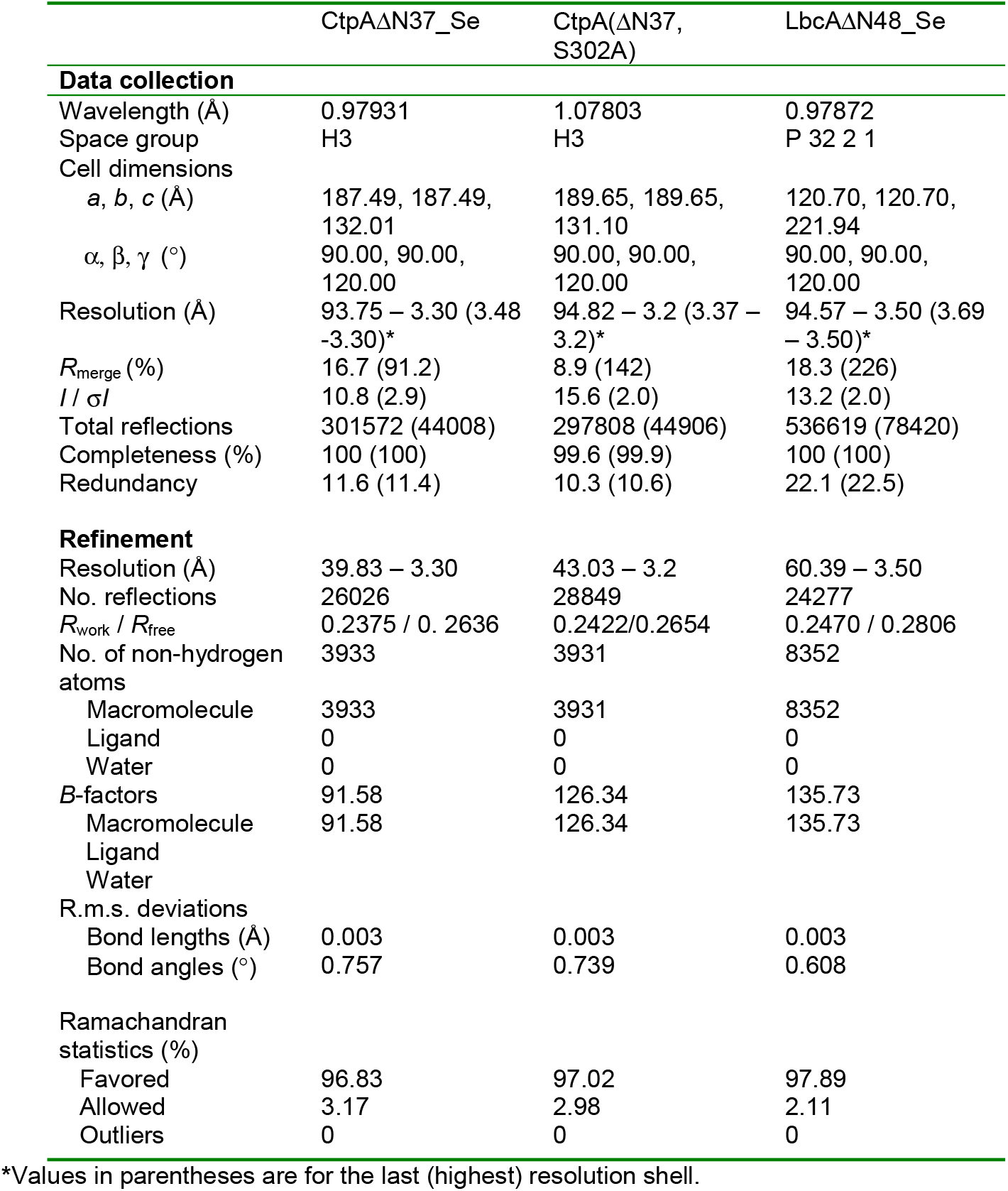
Crystallographic data collection and refinement statistics.

| | CtpAΔN37_Se | CtpA(ΔN37,<br>S302A) | LbcAΔN48_Se |
| --- | --- | --- | --- |
| **Data collection** |  |  |  |
| Wavelength (Å) | 0.97931 | 1.07803 | 0.97872 |
| Space group | H3 | H3 | P 32 2 1 |
| Cell dimensions |  |  |  |
| *a*, *b*, *c* (Å) | 187.49, 187.49,<br>132.01 | 189.65, 189.65,<br>131.10 | 120.70, 120.70,<br>221.94 |
| $\alpha$, $\beta$, $\gamma$ (°) | 90.00, 90.00,<br>120.00 | 90.00, 90.00,<br>120.00 | 90.00, 90.00,<br>120.00 |
| Resolution (Å) | 93.75 – 3.30 (3.48<br>–3.30)* | 94.82 – 3.2 (3.37 –<br>3.2)* | 94.57 – 3.50 (3.69<br>– 3.50)* |
| *R*<sub>merge</sub> (%) | 16.7 (91.2) | 8.9 (142) | 18.3 (226) |
| *I* / $\sigma$ *I* | 10.8 (2.9) | 15.6 (2.0) | 13.2 (2.0) |
| Total reflections | 301572 (44008) | 297808 (44906) | 536619 (78420) |
| Completeness (%) | 100 (100) | 99.6 (99.9) | 100 (100) |
| Redundancy | 11.6 (11.4) | 10.3 (10.6) | 22.1 (22.5) |
| **Refinement** |  |  |  |
| Resolution (Å) | 39.83 – 3.30 | 43.03 – 3.2 | 60.39 – 3.50 |
| No. reflections | 26026 | 28849 | 24277 |
| *R*<sub>work</sub> / *R*<sub>free</sub> | 0.2375 / 0.2636 | 0.2422/0.2654 | 0.2470 / 0.2806 |
| No. of non-hydrogen<br>atoms | 3933 | 3931 | 8352 |
| Macromolecule | 3933 | 3931 | 8352 |
| Ligand | 0 | 0 | 0 |
| Water | 0 | 0 | 0 |
| *B*-factors | 91.58 | 126.34 | 135.73 |
| Macromolecule | 91.58 | 126.34 | 135.73 |
| Ligand |  |  |  |
| Water |  |  |  |
| R.m.s. deviations |  |  |  |
| Bond lengths (Å) | 0.003 | 0.003 | 0.003 |
| Bond angles (°) | 0.757 | 0.739 | 0.608 |
| **Ramachandran**<br>**statistics (%)** |  |  |  |
| Favored | 96.83 | 97.02 | 97.89 |
| Allowed | 3.17 | 2.98 | 2.11 |
| Outliers | 0 | 0 | 0 |

\*Values in parentheses are for the last (highest) resolution shell.

**Supplemental Figure 1.**
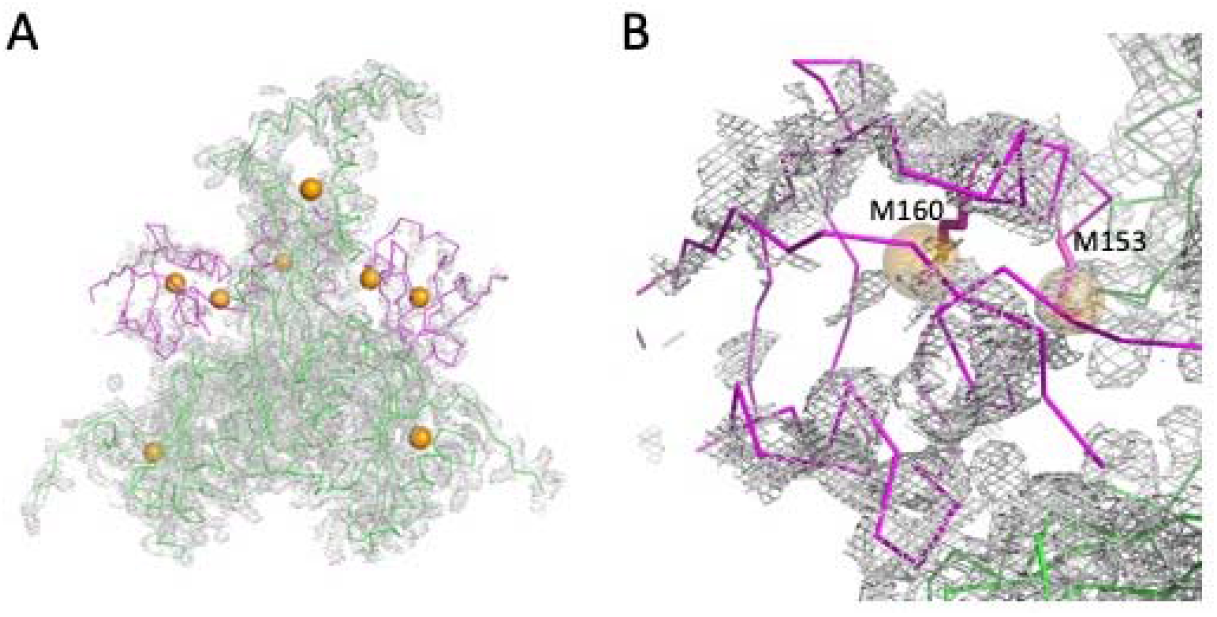
Se-Met peaks superimposed with PDZ homolog model. (A) The overall electron density map of one asymmetric unit obtained from the Se-derivatized CtpA hexamer crystal. The 2mFo-DFc electron density map is rendered at 1σ threshold and shown as gray meshes. The non-PDZ domains are shown in green and the PDZ domains are show in magenta. The positions of Se are in orange. (B) Enlarged view of the left PDZ region in (A).

**Supplemental Figure 2.**
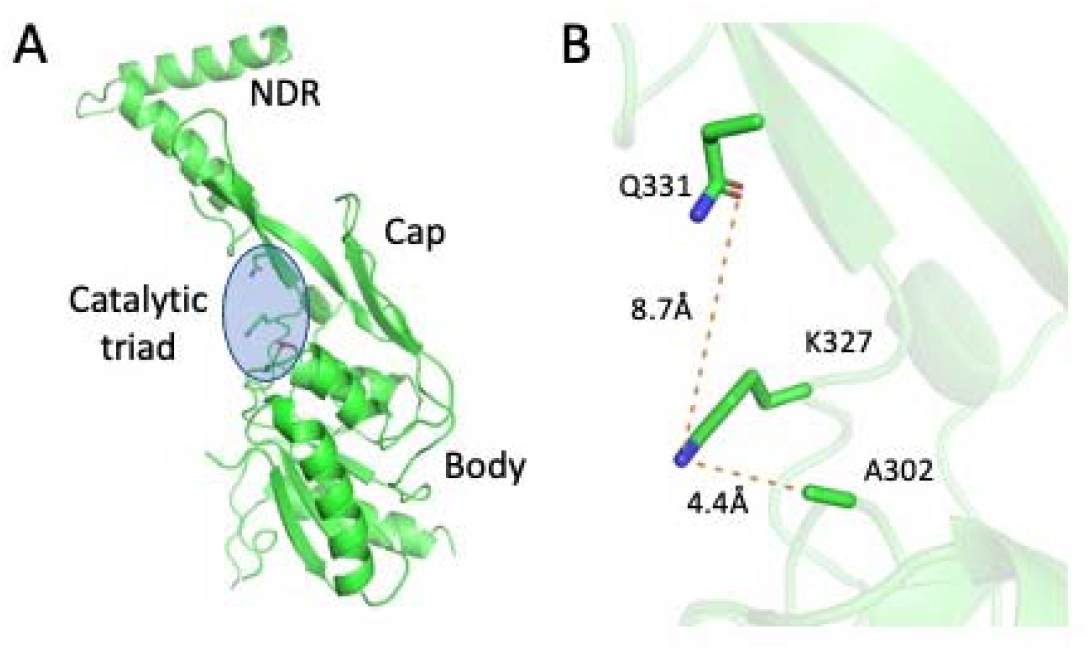
Crystal structure of CtpA(S302A) showing only a monomer. (**A**) overall structure of CtpA(S302A) in ribbons, with the catalytic triad residues in sticks and the point mutation S302A in red sticks. (B) Enlarged view of the catalytic triad in the CtpA(S302A) crystal structure. The structure is also in the inactive configuration.

**Supplemental Video 1. CtpA hexamer structure.** Rotation of the hexamer, then transition to a monomer to show the domain structure, then transition to morphing between inactive and active computational model (based on CtpB).

